# Orphan C/D box snoRNAs modulate fear-related memory processes in mice

**DOI:** 10.1101/2022.05.09.491263

**Authors:** Laura J. Leighton, Qiongyi Zhao, Paul R. Marshall, Sachithrani U. Madugalle, Angelo H. C. Chan, Mason R. B. Musgrove, Haobin Ren, Ambika Periyakaruppiah, Janette Edson, Wei Wei, Xiang Li, Robert C. Spitale, Timothy W. Bredy

## Abstract

C/D box small nucleolar RNAs (snoRNAs) comprise a class of small noncoding RNAs with important regulatory effects on cellular RNA function. Although it is well established that snoRNAs coordinate the post-transcriptional modification of pre-ribosomal and small nuclear RNAs by 2’-O-methylation, which leads to enhanced RNA stability, whether they are necessary for memory-related processes remains relatively unexplored. Using targeted sequencing, we have identified more than 150 C/D box snoRNAs in the prefrontal cortex of male C57BL/6J mice, 31 of which are differentially expressed in response to fear extinction learning. We have also discovered a subset of snoRNAs, including many orphans, that are enriched in the synaptic compartment, including the orphan snoRNA snord64. An extinction learning-induced increase in synapse-enriched snord64 led to increased 2’-O-methylation within the 3-UTR of the mRNA encoding the ubiquitin ligase RNF146. This effect was blocked by snord64 knockdown and was accompanied by attenuated forgetting of conditioned fear and the enhanced retrieval of fear extinction memory. Localized activity of orphan snoRNAs therefore represents a novel mechanism associated with fear-related learning and memory.

**Significance statement:** We have discovered a population of experience-dependent small nucleolar RNAs (snoRNAs), and found that several of these are recruited to the synaptic compartment in response to fear extinction learning. In particular, the orphan snoRNA, snord64, drives the methylation of the mRNA encoding the ubiquitin ligase, RNF146, with a reduction in snord64 attenuating forgetting of conditioned fear and enhancing the retrieval of fear extinction memory. This study reveals a new mechanism of gene regulation associated with fear-related memory that involves the activity of synapse-enriched snoRNAs.

## Introduction

RNA possesses additional layers of information-carrying capacity beyond its nucleotide sequence and its abundance within the cell. Secondary structure and covalent modification of RNA is critical for its function, at times acting as an environmentally responsive switch to alter the function of individual RNA molecules in a context- and state-dependent manner (reviewed by Wan et al., 2011; Mortimer et al., 2014; Bevilacqua et al., 2016). The importance of these features is especially clear in neurons, which transport RNA to synapses physically distant from the cell body, and can undertake activity-dependent local translation of synaptic RNA (Biever et al., 2019). Identification of neuron-specific RNA modifications and their functions is therefore critical to understanding rapid local responses to incoming information.

Many RNA modifications are driven by proteins which recognize sequence or structural elements within target RNAs. Other modifications are guided by small RNAs which act as adapters between a modifying complex and the target RNA through base pairing. This mechanism is exemplified by small nucleolar RNAs (snoRNAs), of which there are two classes, each defined by specific sequence motifs. H/ACA box snoRNAs guide the isomerization of specific uridine residues to pseudouridine, whereas C/D box snoRNAs direct 2’-O-methylation of complementary RNAs, thereby impacting their stability and secondary structure (Figure 1) (Reichow et al., 2007; Henras et al., 2017). The majority of snoRNAs guide the modification of pre-ribosomal RNA, with these snoRNA-dependent rRNA modifications being highly conserved, necessary for the structure of the ribosome, and essential for survival (Reichow et al., 2007; Henras et al., 2017). However, a substantial number of snoRNAs are orphans, and have no identified complementarity to ribosomal sequences. The function of orphan snoRNAs remains largely unknown, although prior studies have found that some of them exhibit brain-specific patterns of expression (Cavaillé et al., 2000; Runte et al., 2001; Rogelj, 2006). Beyond this, little is known about the role of orphan snoRNAs in the brain or within the context of learning and memory.

**Figure 1:**
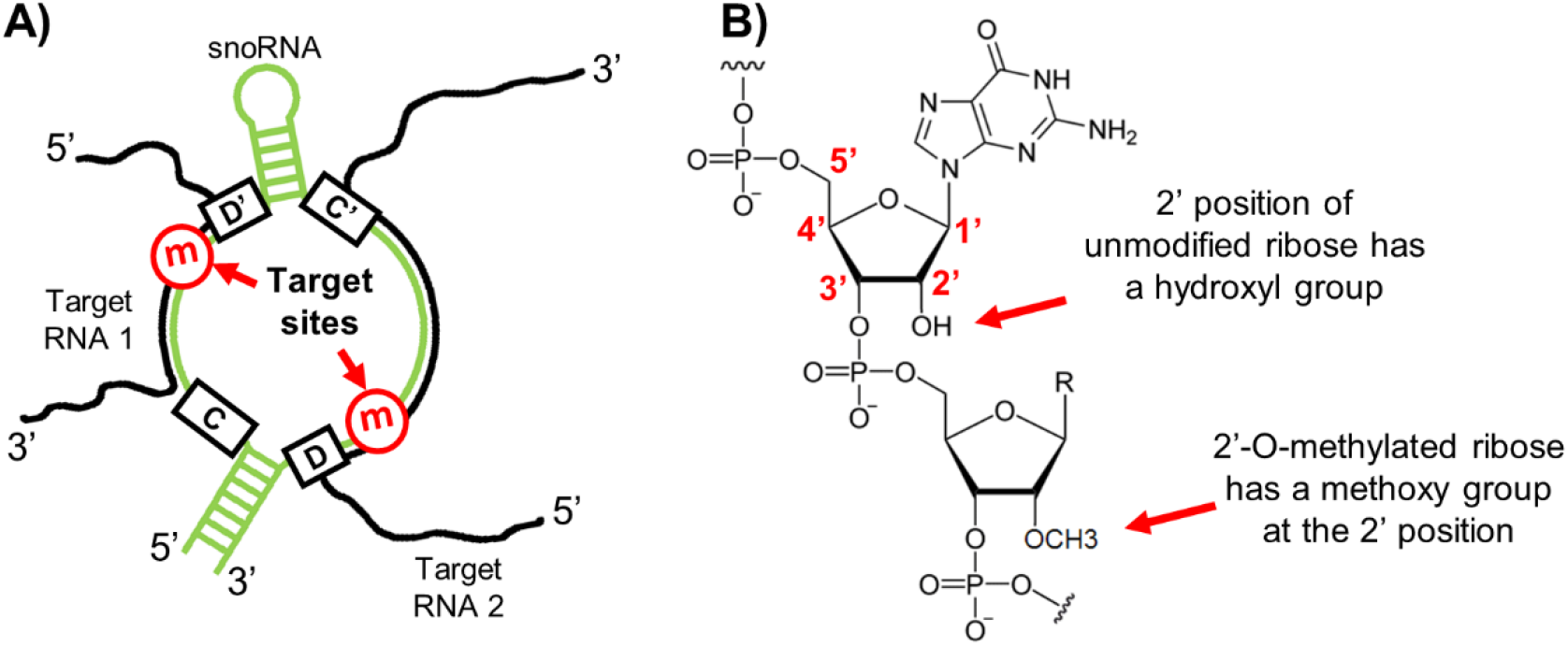
Structure and function of C/D box snoRNAs. C/D box snoRNAs (A) fold into a single loop structure which has two open regions which base-pair with targets to guide 2’-O-methylation. This modification (B) occurs on the ribose group of the RNA backbone,and can co-occur with any nucleobase.

Learning to respond fearfully to stimuli which are associated with negative events is a broadly conserved survival mechanism, as is the extinction of the fear response following the repeated presentation of a feared cue with no further negative consequences. Our group and others have demonstrated the importance of activity-dependent gene regulation for the learning, recall and extinction of conditioned fear (Lin et al., 2011; Baker-Andresen et al., 2013; Katz and Lamprecht, 2015; Peixoto et al., 2015; Leighton et al., 2019; Li et al., 2019; Mizuno et al., 2020; Wei et al., 2022). Given that few studies have considered whether C/D box snoRNAs are involved in memory related processes, we set out to address this by examining the experience-dependent expression of C/D box snoRNAs in the adult mouse medial prefrontal cortex (mPFC). We identified over 150 snoRNAs, including 31 snoRNAs that show dynamic activity in response to fear extinction learning, a substantial number of which are orphans. Further investigation demonstrated that one such snoRNA, snord64, dynamically enters the synaptic compartment and guides the learning-induced 2’-O-methylation of the mRNA encoding a neuroprotective ubiquitin ligase, *Rnf146*. Knockdown of snord64 attenuated the forgetting of conditioned fear and enhanced the retrieval of fear extinction, suggesting localised snoRNA activity is required for memory related processes.

## Materials and Methods

### Animals

All experiments involving animals were approved by the Animal Ethics Committee of The University of Queensland. Male C57BL/6J mice (10-20 weeks old) were used for all behavioral experiments. Mice were housed two per cage separated by a clear plastic divider and were provided with nesting materials and access to food and water ad libitum. Mice were maintained on a 12-hour light/dark cycle, with all behavioral experiments being carried out during the light phase.

### Fear conditioning and extinction

Mice underwent fear conditioning in Context A (square shape, medium size, yellow lighting, wire grid floor, lemon scent). The fear conditioning protocol consisted of 3 presentations of a 2-minute pure tone conditioned stimulus (CS) paired with a 1-second, 0.7mA foot shock. The animals used for the snoRNA sequencing experiment underwent trace fear conditioning (foot shock 10 seconds after the end of the CS). All other animals underwent delay fear conditioning (foot shock co-terminating with the CS.)

Animals used for tissue sampling for molecular experiments were randomly assigned to extinction or retention-control groups. On the day following fear conditioning, mice in the extinction group were exposed to 60 presentations of the CS in Context B (round shape, small size, green lighting, plastic floor, vinegar scent.) Mice in the retention control group were placed in Context B for the same length of time, but not exposed to the CS. Immediately after extinction or retention control training, mice were euthanized by cervical dislocation and the PFC was rapidly dissected on a chilled metal platform, then snap-frozen on dry ice.

### Sequencing of snoRNAs from the mouse PFC

Nuclei were isolated from frozen mouse PFC samples according to a published protocol (Jiang et al., 2008; Li et al., 2014). Briefly, whole mouse PFC was mechanically dissociated using a glass douncer, after which dissociated cells were lysed in hypotonic nuclear extraction buffer and nuclei collected by gentle centrifugation. RNA was isolated from cell nuclei using the NucleoSpin miRNA kit (Macherey-Nagel), following the manufacturer’s directions to isolate small RNA and large RNA in separate fractions. Sequencing libraries were prepared from 50ng nuclear small RNA using the NEBNext Small RNA Library Prep Set for Illumina (New England Biolabs). Libraries were pooled and run on 10% TBE/polyacrylamide gels at 60 volts for 8-12 hours at 4°C, and the region from approximately 175-235bp (corresponding to an insert size of 50-110nt) was cut from the gel and DNA isolated using the crush-and-soak method. Single-end sequencing was performed on the Illumina Hiseq 2000 using V4 chemistry and a read length of 1×126nt. Image processing and basecalling were performed using the standard Illumina Genome Analyzer software. Adapter sequences were trimmed using Cutadapt (v1.8.1) and alignment to the mouse reference genome (mm10) was performed using Bowtie (v1.2.1.1) with the option “--best --strata --v 1” to report only those alignments in the best alignment ‘stratum’ with at most one mismatch. Custom PERL scripts were applied to generate the raw count and CPM (counts per million) table. Manual deconvolution of the dataset was performed by collating sequences overlapping at a single locus under a single identifier. Manual annotation of C/D box snoRNAs was performed based on the presence of C and D motifs, and identification of snoRNAs by searching for the most abundant sequence representing each unique small RNA in the snoRNA Orthological Gene Database (snOPY) (Yoshihama et al., 2013), and when necessary, other online databases and the literature. Differential expression analysis was performed on the collated list of 164 CD box snoRNAs using GraphPad Prism to perform multiple heteroscedastic t-tests, and discovery was determined using the two-stage linear step-up procedure of Benjamini, Krieger and Yekutieli, with a false discovery rate (Q) of 5%.

### Synaptosome preparations

The synaptosome isolation protocol was adapted from previously published protocols (Dunkley et al., 2008; Westmark et al., 2011). Four mouse PFCs per sample from the same behavioral group were gently homogenized in 2mL of gradient medium buffer supplemented with 1µL/mL RNaseOUT and 1mM dTT. Samples were centrifuged at 1000 x *g* for 10 minutes at 4°C to pellet the non-synaptic fraction, which was snap-frozen. The supernatant was loaded onto a discontinuous density gradient consisting of layers of 23%, 15% and 3% Percoll diluted with iso-osmotic sucrose. Samples were separated by ultacentrifugation for 5 minutes at 30,700 x *g* and 4°C, and the synaptosome fraction was isolated by gentle pipetting. Synaptosomes were washed once with gradient medium buffer and collected by ultracentrifugation for 30 minutes at 19,900 x *g* and 4°C. The recovered synaptosomes were washed with PBS and pelleted in a benchtop centrifuge, then snap-frozen. To isolate RNA, samples were homogenized in Nucleozol (Macherey-Nagel); water was added and samples were centrifuged to precipitate contaminants according to the manufacturer’s directions, after which the supernatant was mixed 1:1 with ethanol and RNA captured on a Zymo-Spin IC column (Zymo Research). DNase treatment was performed on-column and RNA was washed and eluted according to the manufacturer’s directions. RNA sequencing libraries were prepared from the synaptosomal and non-synaptic fractions of 6 synaptosome preps (3 from 60CS fear extinction and 3 from time-equivalent retention controls) using the SMARTer Stranded Total RNA-seq kit v2 - Pico Input Mammalian (Takara Bio). Paired-end sequencing with a read length of 2×75 was performed on the Illumina Hiseq 4000 by Genewiz.Cutadapt (v1.17) was used to clip low-quality bases and adapter sequences, and processed reads were aligned to the mouse genome (mm10) using HISAT2 (v2.1.0). HTSeq-count (v0.11.0) was used to generate the raw count table for each sample. EdgeR (v3.26.8) was used for differential expression analysis.

### Reverse transcription and quantitative PCR

For quantitative PCR of mRNA targets, 100-500ng of high-quality total RNA per reaction was reverse transcribed using the QuantiTect RT kit (Qiagen) according to the manufacturer’s instructions, using the provided RT Primer Mix. cDNA was diluted at least 1:1 with water.

For quantitative PCR of snoRNA targets, a ligation reaction was prepared from 100ng of high-quality total RNA, 5 picomoles of 5’-adenylated and 3’-blocked linker, 200 units of T4 RNA ligase 2, truncated KQ, and 20% polyethylene glycol (PEG8000). Ligation reactions were incubated at 25°C for 2 hours, after which a reverse transcription reaction mix containing Protoscript II reverse transcriptase and a primer complementary to the ligated linker was added. Reverse transcription was performed at 50°C for 1 hour, and the reactions were then diluted 1:1 with water.

Quantitative PCR reactions were prepared in duplicate, in a 10µL reaction volume, using 2X Sensifast SYBR green master mix (Bioline), 500µM of each primer, and 1µL per reaction of cDNA sample. Reactions were run on the Rotor-Gene Q platform and results were analyzed using the delta-delta-cT method. Normalization was performed to *Pgk1* (phosphoglycerate kinase) for mRNA targets, and 5.8S rRNA for snoRNA targets.

### Reverse Transcription at Low dNTP concentrations followed by PCR (RTL-P)

For site-specific assay of internal mRNA 2’-O-methylation, cDNA synthesis was performed in duplicate for each sample, using 30-100ng of high-quality total RNA and a pool of experiment-dependent gene-specific primers. One reverse transcription reaction per sample contained a standard quantity of dNTPs (final concentration 0.5mM) while the other reaction contained one hundredfold less (final concentration 5µM). Reverse transcription was performed using SuperScript III according to the manufacturer’s directions. For each of the two cDNA preparations per sample (high-dNTP and low-dNTP), quantitative PCR was performed with each of two primer sets, one located immediately upstream of the putative 2’-O-Me site, and one located immediately downstream. Quantitative PCR reactions were prepared and run as described previously. Data were analyzed using the delta-delta-cT method, normalizing the upstream primer set to the downstream primer set for each sample to determine the readthrough percentage, *r* (corresponding to unmethylated transcripts). Methylation level was calculated as 100 - *r*.

### Design of antisense oligonucleotides for snoRNA knockdown

Two antisense oligonucleotides (ASOs) were designed against regions of snord64 predicted to be single-stranded by Mxfold (Akiyama et al., 2018), one of which approximately corresponds to the predicted antisense element. The sequence of the negative control ASO was designed by Integrated DNA Technologies, bioinformatically predicted to lack targets in either human or mouse, and has been validated by IDT for use in knockdown experiments (Lennox and Behlke, 2020). All ASOs were 20 nucleotides long with a total phosphorothioate backbone, and used a 5-10-5 gapmer design with 5 nucleotides at either end modified with 2’-O-methoxyethyl (2’MOE). Upon receipt, ASOs were resuspended in ultrapure water at 200µM concentration and stored at -20°C.

### Stereotaxic surgery for cannula placement

Male C57BL/6J mice (10-12 weeks old) were anaesthetized by intraperitoneal injection of 100mg/kg ketamine and 10mg/kg xylazine. The surgical site was shaved and disinfected with iodine followed by 80% ethanol, and the mouse was then secured in a stereotaxic frame. A midline skin flap was removed using surgical scissors and the periosteum was removed using 80% ethanol. A burr hole was opened over the prefrontal cortex and a steel double cannula was lowered into the infralimbic prefrontal cortex at coordinates ML visual midline, AP +1.8mm, DV-2.7mm relative to bregma. Cannulae were permanently anchored using acrylic dental cement with one supporting screw in each parietal plate. At the completion of surgery, analgesia was provided by subcutaneous injection of buprenorphine and meloxicam, repeated if required for up to 3 days. Mice were allowed to recover from surgery for a minimum of one week before commencement of behavioral experiments.

### Intracerebral ASO infusion and behavioral tests

After recovery from surgery, mice underwent delay fear conditioning as described above. Following fear conditioning, mice with a 3^rd^-CS freezing score below 20% were excluded from the experiment, and remaining mice were assigned into four experimental groups so that all groups had equal numbers of mice and average 3^rd^-CS freezing scores that were as similar as possible. 24 hours after fear conditioning, mice were infused with either the negative control ASO, or a 1:1 mixture of snord64-ASO1 and snord64-ASO2. ASOs were prepared for injection using the *in vivo-*jetPEI delivery reagent (Polyplus) according to the manufacturer’s instructions. A total of 500ng of ASO was used for each animal, in a total injection volume of 2µL isotonic glucose with an N/P ratio of 6. Mice were lightly anaesthetized with isoflurane and 1µL ASO was infused through each side of the cannula over a 5 minute period using a Hamilton syringe connected to a syringe driver. Following completion of the injection, the injector was left in place for 30-60 seconds. One week after ASO infusion, mice in the extinction groups were exposed to 10 presentations of the CS in Context B, while mice in the retention control groups were placed in Context B for the same length of time without presentation of the CS. Recall tests were performed according to the schedule shown in Figure 6E. Analysis of freezing behaviour was performed using FreezeFrame 5 software (Actimetrics) by a researcher blind to experimental group.

## Results

### Learning-induced expression of snoRNAs in the medial prefrontal cortex

To explore the abundance and diversity of experience-dependent C/D box snoRNA expression, we deeply sequenced small RNAs from the PFC of 5 male mice following fear extinction learning, and 5 retention controls. Samples were expected to represent all cell types of the PFC, and were enriched for nuclear RNAs of approximately 50-110 nucleotides (Figure 2A). 164 small RNAs passing an abundance filter (minimum 1 read per million in all five samples from one experimental group) and possessing the C-box and D-box motifs which define C/D box snoRNAs were considered for further analysis.

**Figure 2:**
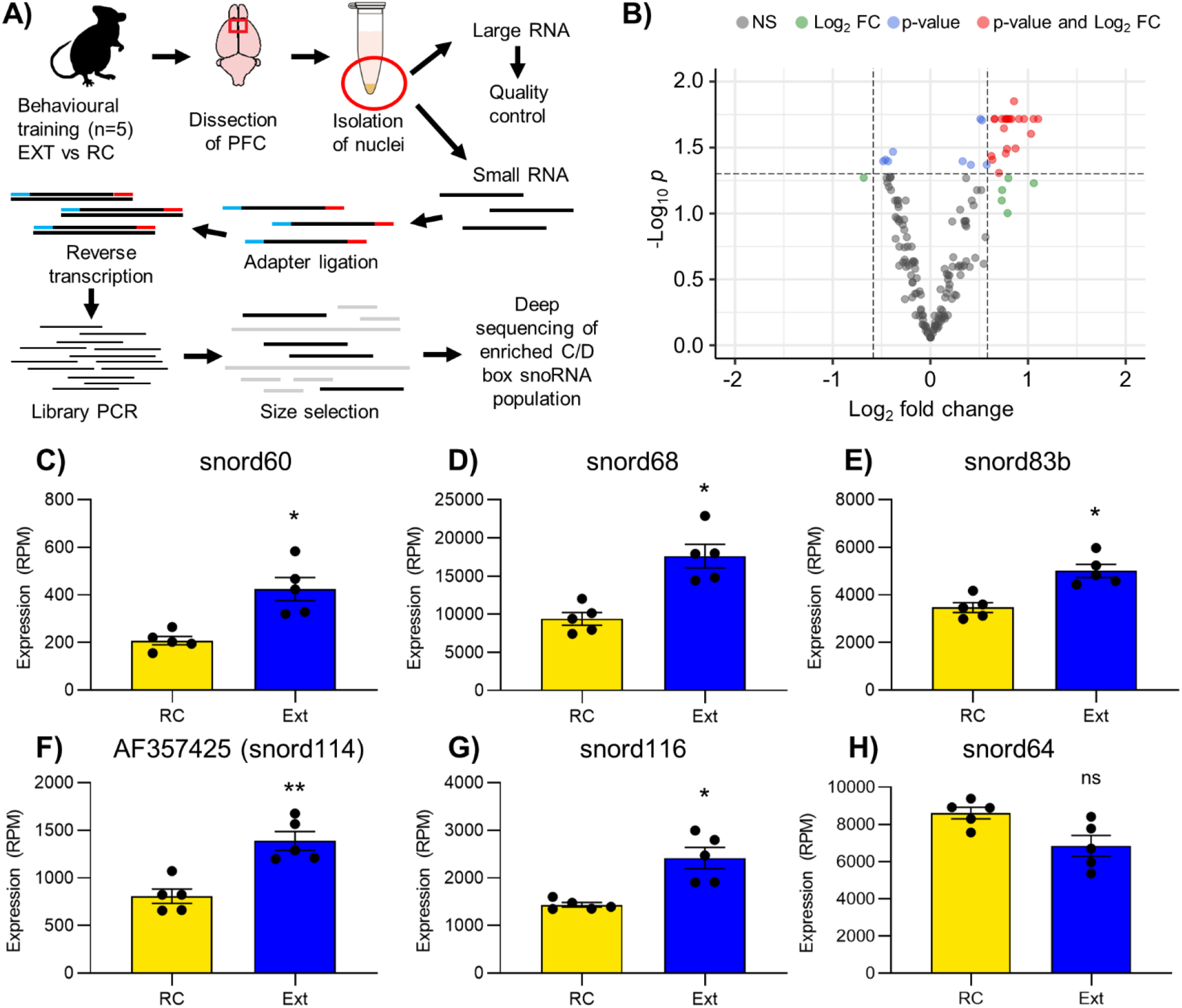
Activity dependent expression of snoRNAs from the mouse PFC. A) Overview of sample preparation for enrichment and sequencing of *CID* box snoRNAs. 8) Volcano plot of snoRNAs showing differential expression of some in fear extinction learning relative to retention control. C-H) Activity dependent changes in the expression of 6 snoRNAs within the PFC in response to fear extinction learning relative to retention controls. C) snord60 is upregulated in Ext (t-test, p=0.003046, q=0.024878.) D) snord68 is upregulated in Ext (t-test, p=0.001439, q=0.019201.) E) snord83b is upregulated in Ext (t-test, p=0.002139, q=0.019656.) F) AF37425, an instance of snord114, is upregulated in Ext(t-test, p=0.001541, q=0.0019201.) G) snord116 is upregulated in Ext (t-test, p=0.002615,q=0.022613.)H) snord64 trends downwards in Ext (p=0.025447, q=0.07959.)

To identify the snoRNAs, we used the snoRNA database snOPY (Yoshihama et al., 2013), with additional information obtained from snoRNA-LBME-db (Lestrade and Weber, 2006), Rfam and the UCSC genome browser. The snOPY database contains records for 327 mouse C/D box snoRNAs: 177 records representing multiple instances of the tandemly repeated snoRNAs snord113/114, snord115 and snord116, and 150 records representing non-repeat-associated snoRNAs. We detected expression of some copies of each repeat-associated snoRNA, and 127 non-repeat-associated snoRNAs, representing 84.7% of well-annotated mouse C/D box snoRNAs. We also detected 33 small RNAs with C/D box motifs which did not have associated records in the snOPY database. Of these, 29 were recognized as C/D box snoRNAs by at least one of snoRNA-LBME-db, Rfam, RNAcentral or the UCSC genome browser and a further two showed both sequence homology and synteny with recognised human C/D box snoRNAs.

Most C/D box snoRNAs have known target sites on ribosomal RNA. However, a significant minority are orphans, with no complementarity to ribosomal RNA and no known targets. When characterizing orphan snoRNAs, we made the assumptions that mouse homologs of human snoRNAs with known target sites were not orphans, and that snoRNAs so poorly annotated that they lacked records in both snOPY and snoRNA-LBME-db were orphans. We found that 35 of 164 C/D box snoRNAs (21.3%) were orphans.

All of the C/D box snoRNAs identified from the mouse PFC were detected in all samples from both behavioral groups. 31 snoRNAs (18.9%) were differentially expressed in the nucleus between mice which underwent fear extinction learning and retention controls. Of these, 27 were upregulated while 4 were downregulated (Table 1; Figure 2B). The average fold change for differentially expressed snoRNAs was small (0.638).

Given that most C/D box snoRNAs are hosted within introns of mRNAs and lncRNAs, we next asked whether differential expression of snoRNAs following fear extinction learning was explained by altered expression of their host transcripts. Because many snoRNA host genes contain multiple introns with snoRNAs, the 31 differentially expressed snoRNAs are hosted within 25 different transcripts. We examined expression of these 25 transcripts in RNA sequencing data derived from the non-synaptic fraction of mouse PFC lysates. None of the 25 snoRNA host genes was differentially expressed in response to fear extinction learning, indicating that differential regulation of the hosted snoRNAs in response to behavioral experience is independent of regulation of the host transcripts.

### C/D box snoRNAs are present at within the synaptic compartment in the medial prefrontal cortex

To identify RNAs which are localized to the synaptic compartment of the mouse PFC, RNA was sequenced from synaptosomes and from the non-synaptic fraction of mouse prefrontal cortex homogenate. There was no detectable contamination of the synaptosomal fraction with nuclear material. Unexpectedly, some snoRNAs were detected both in the nonsynaptic fraction (which is enriched for nuclei and perinuclear soma) and in synaptosomes, including a striking number of orphan snoRNAs.

We validated the presence of 6 C/D box snoRNAs at synapses by qRT-PCR in a different set of synaptosomal and non-synaptic RNA samples, comparing extinction and retention control. Snord8 and snord67 were present at the synapse, and were stably expressed between fear extinction and retention control in both synaptic and non-synaptic fractions (Figure 3A, B.) Snord22 and snord83b increased in abundance at the synapse in fear extinction relative to retention controls (Figure 3C, D.) The most interesting pattern was observed for snord99 and snord64, both of which decreased in abundance in the non-synaptic fraction but increased in abundance at the synapse during fear extinction (Figure 3E, F.)

**Figure 3:**
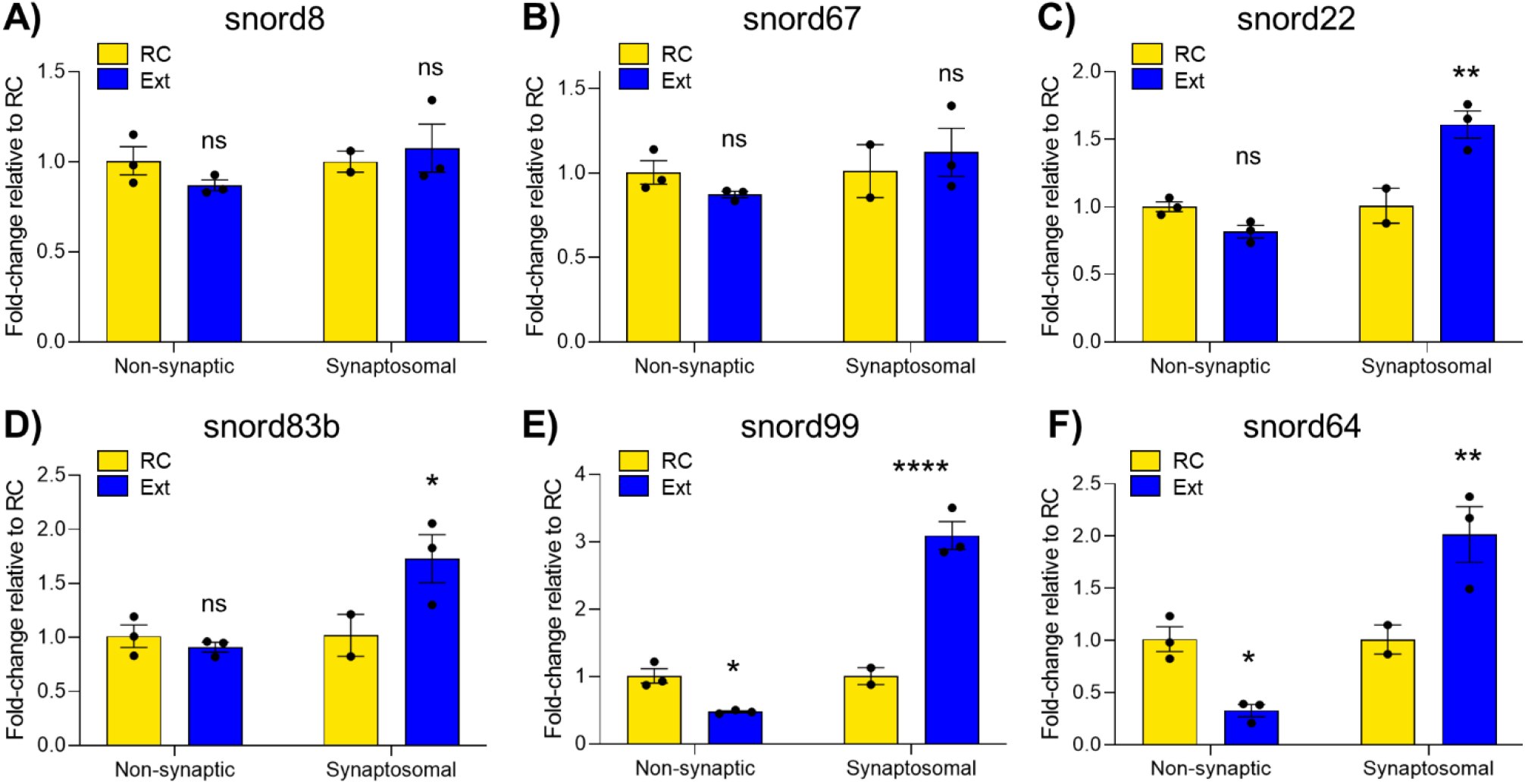
Patterns of localisation between extinction learning and retention control for six snoRNAs with substantial expression at the synapse. A) snord8 shows no difference between EXT and RC in the non-synaptic fraction (two-way ANOVA, no significant effects, n=3, Sidak’s multiple comparisons test p=0.5102) or at the synapse (Sidak p=0.8415.) B) snord67 shows no difference between EXT and RC in the non-synaptic fraction (two-way ANOVA, no significant effects, n=3, Sidak’s multiple comparisons test p=0.6088) or at the synapse (Sidak p=0.7482.) C) snord22 shows no difference between EXT and RC in the non-synaptic fraction (two-way ANOVA, significant main effect of compartment p=0.0014, significant main effect of behaviour p=0.0319, significant interaction p=0.0015, Sidak’s multiple comparisons test p=0.2251) but is significantly increased at the synapse in EXT compared to RC (Sidak p=0.0026). D) snord83b shows no difference between EXT and RC in the non-synaptic fraction (two-way ANOVA, significant main effect of compartment p=0.0354, no significant main effect of behaviour p=0.0918, significant interaction p=0.00354, Sidak’s multiple comparisons test p=0.8717) but is significantly increased at the synapse in EXT compared to RC (Sidak p=0.0366). E) snord99 decreases in the non-synaptic fraction (two-way ANOVA, significant main effect of compartment p<0.0001, significant main effect of behaviour p=0.0008, significant interaction p<0.0001, Sidak’s multiple comparisons test p=0.0464) and increases at the synapse (Sidak p<0.0001). F) snord64 decreases in the non-synaptic fraction (two-way ANOVA, significant main effect of compartment p=0.0020, no significant main effect of behaviour p=0.3895, significant interaction p=0.0019, Sidak’s multiple comparisons test p=0.0435) and increases at the synapse (Sidak p=0.0125.) *p<0.05,**p<0.01, ****p<0.00001.

It is important to note that, for each snoRNA, relative expression analysis was performed separately for the non-synaptic and synaptosomal fractions due to the non-availability of a reference small RNA which is equally distributed throughout all subcellular compartments. The scaling of the graphs in Figure 3 does not indicate that baseline expression of each snoRNA in the synaptosomal and non-synaptic fractions is the same. For all six snoRNAs measured, the raw cT values for the non-synaptic fraction were 4-6 cycles lower than for the synaptosomal fraction, indicating that each snoRNA is considerably more abundant in the nucleus or perinuclear soma than at the synapse.

### C/D box snoRNAs guide site-specific 2’-O-methylation of mRNAs in the medial prefrontal cortex

To investigate the hypothesis that orphan C/D box snoRNAs have non-ribosomal targets in the mouse PFC, we selected the candidate gene snord64. This snoRNA is encoded within the Prader-Willi syndrome critical region, a paternally expressed imprinted domain which contains numerous C/D box snoRNAs. Prader Willi syndrome, caused by loss of expression of this domain, is characterized by mild intellectual disability, hyperphagia and obesity (Angulo et al., 2015). Expression of snord64 in nuclear small RNA sequencing libraries trended downwards in extinction-trained mice relative to retention controls, although this effect did not reach statistical significance with multiple comparisons correction applied (p=0.024557, q=0.0795, Figure 2H.) However, this small RNA was detected in synaptosomes, and its synaptic abundance was significantly increased in extinction-trained mice relative to retention controls (Figure 3F), which may indicate that it is trafficked to the synaptic compartment during learning. Bioinformatic prediction of potential snoRNA:mRNA interactions using PLEXY (Kehr et al., 2011) identified a possible interaction between snord64 and the 3’ UTR of the messenger RNA encoding Ring finger nuclease 146 (*Rnf146*), a neuroprotective ubiquitin ligase (Figure 4B). Although snord64 is an orphan, it has intact C and D motifs, and meets the structural requirements for a methylation guide as described by Qu *et al* (Qu et al., 2011). The predicted interaction between snord64 and *Rnf146* is 19 nucleotides long and contains 14 canonical base pairs, 5 G:U base pairs, and no mismatches; this interaction is stronger than that between snord32a/snord51 and the peroxidasin (*Pxdn*) mRNA which is confirmed to guide methylation (Elliott et al., 2019). We found that *Rnf146* is methylated in the mouse PFC (Figure 4C). The methylation level increases from 60% in retention control mice to 74% following fear extinction learning (Figure 4D). This finding demonstrates that the novel 2’-O-methylation site on *Rnf146* is a dynamic epitranscriptomic mark that undergoes a rapid learning-induced response, *in vivo*.

**Figure 4:**
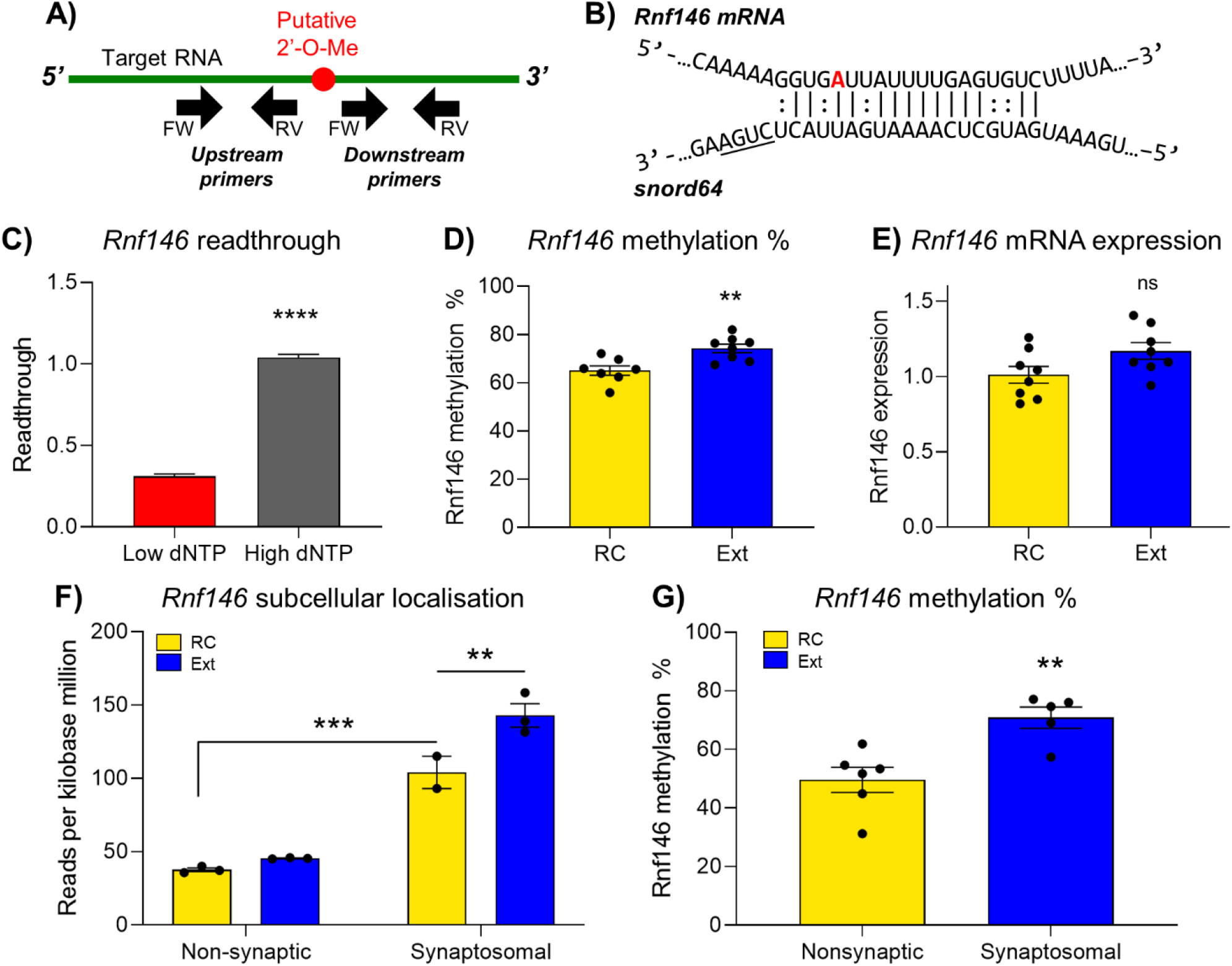
snord64 methylates *Rnf146* mRNA and transcripts with this modification show dynamic subcellular localisation changes. A) Overview of RTL-P, a qRT-PCR based assay for quantitative analysis of 2’-O-methylation. B) snord64 was predicted to interact with *Rnf146* with imperfect complementarity through a 17-nucleotide antisense region which contains 14 canonical base pairs, five G:U base pairs and no mismatches. C) Readthrough of *Rnf146* mRNA (upstream primer set normalised to downstream primer set) is reduced in the mouse prefrontal cortex under low dNTP conditions (n=16) compared to high dNTP conditions (n=16) (Student’s t-test, p<0.0001), indicating that the transcript is methylated at the predicted snord64 interaction site. D) The methylation level of *Rnf146* in the mouse PFC increases in fear extinction (n=8) relative to retention controls (n=8) (Mann-Whitney test, p=0.0030.) E) There is no significant difference in *Rnf146* mRNA expression between extinction (n=8) and retention control (n=8) (Student’s t-test, p=0.0636.) F) Data from RNA sequencing comparing synaptosomal to non-synaptic samples shows that the baseline abundance of *Rnf146* is greater at the synapse (two-way ANOVA, n=3, significant main effect of compartment p<0.0001, significant main effect of behaviour p=0.0052, significant interaction p=0.0326, Sidak’s multiple comparisons test p=0.0002.) The abundance of *Rnf146* at the synapse is significantly increased in fear extinction compared to equivalent retention control (two-way ANOVA, n=3, significant main effect of compartment p<0.0001, significant main effect of behaviour p=0.0052, significant interaction p=0.0326, Sidak’s multiple comparisons test p=0.0058.) G) The methylation level of *Rnf146* at the snord64 interaction site is greater at the synapse than in the non-synaptic fraction (Student’s t-test, n=5-6, p=0.0051.) **p<0.01, ****p<0.0001.

Given that snord64 abundance declines in the non-synaptic fraction but increases at the synapse during fear extinction learning (Figure 3F), we next asked where its methylation target *Rnf146* was located in the cell. RNA sequencing from synaptosomes showed that *Rnf146* is approximately three times more abundant at the synapse than in the non-synaptic fraction (Figure 4F). In addition, the abundance of *Rnf146* at the synapse increased in response to fear extinction learning, while there was no change in the quantity of this transcript in the non-synaptic fraction (Figure 4F) or in whole mouse PFC samples as measured by quantitative PCR (Figure 4E). We also found that the proportion of *Rnf146* transcripts which are methylated at the snord64 interaction site is greater at synapses than in the nonsynaptic compartment (Figure 4G).

We also investigated the C/D box snoRNA snord115, which is encoded in tandemly repeated clusters within the Prader-Willi syndrome critical region, and which was the first reported example of snoRNA:mRNA complementarity (Cavaillé et al., 2000). Snord115 interacts with the messenger RNA encoding the serotonin receptor 2C (*Htr2c*) through an 18-nucleotide targeting element with perfect complementarity (Figure 5A.) While this interaction is known to influence splicing and adenosine-to-inosine editing of this mRNA (Vitali et al., 2005; Kishore and Stamm, 2006), studies which detailed this interaction in depth did not include experiments to determine whether *Htr2c* was 2’-O-methylated, while some authors have argued that it is not (Kishore et al., 2010; Bratkovič et al., 2018). We found that *Htr2c* is highly methylated in the mouse prefrontal cortex (Figure 5B). The methylation level is stable between extinction and retention control (Figure 5C), suggesting that the constitutive state of this transcript is to be heavily methylated. This finding has relevance to the role of snord115 in Prader-Willi syndrome and validates the use of *Htr2c* as a positive control for mRNA 2’-O-methylation in mouse brain samples.

**Figure 5:**
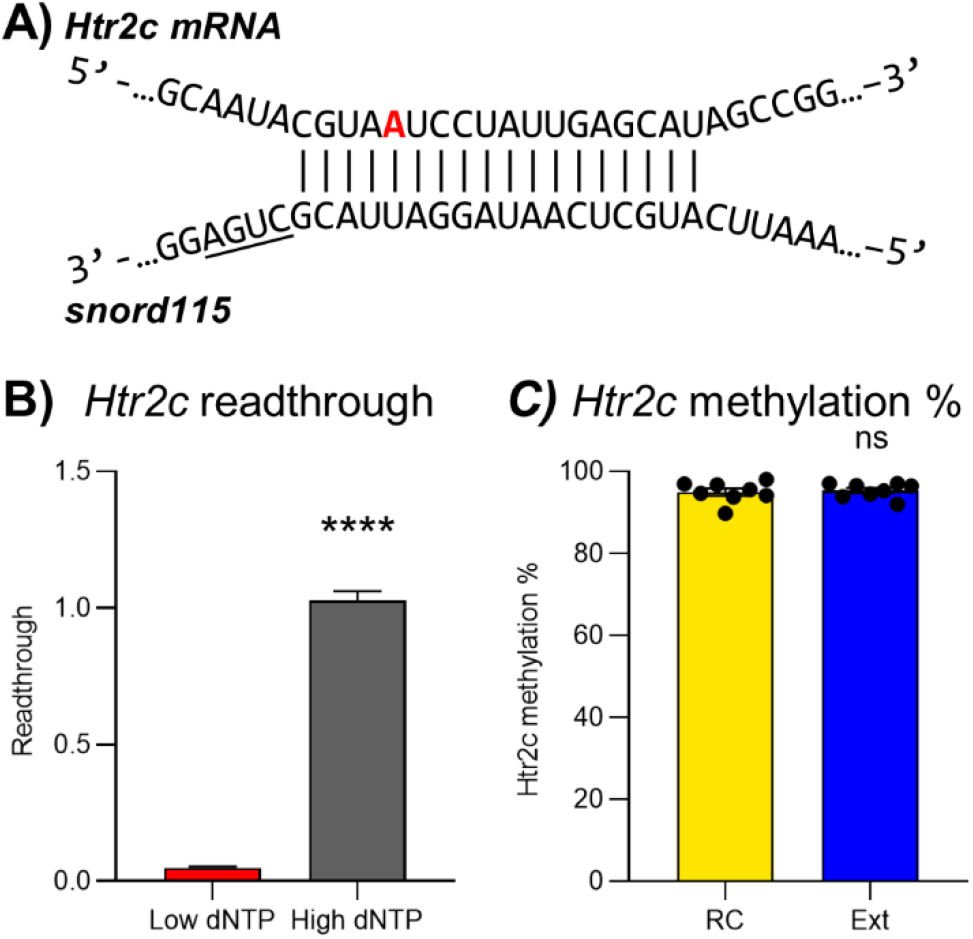
*Htr2c* is methylated in the mouse PFC. A) snord115 interacts with *Htr2c* mRNA with perfect complementarity through an 18-nucleotide antisense region. B) *Htr2c* readthrough is dramatically reduced in the mouse prefrontal cortex when reverse transcription is performed under low dNTP conditions (n=16) compared to high dNTP conditions (n=16) (Student’s t-test, p<0.0001), indicating near-complete methylationof this transcript. There is no difference in the methylation level of *Htr2c* between fear extinction (n=8) and retention control (n=8) (Student’s t-test,p=0.7528).

### Knockdown of snord64 enhances the retention of fear-related memory

To reduce snord64 expression, two antisense oligonucleotides (ASOs) were designed against predicted structurally open regions of snord64, one of which approximately corresponded with the antisense element targeting *Rnf146* (Figure 6A). The ASOs were 20-nucleotide, phosphorothioate-stabilized 2’MOE gapmers; this RNase H-dependent design was chosen because it has previously been successfully used to knock down snoRNAs and importantly, knockdown of snoRNAs using ASOs does not affect the abundance of the host transcript (Liang et al., 2011; Michel et al., 2011).

**Figure 6:**
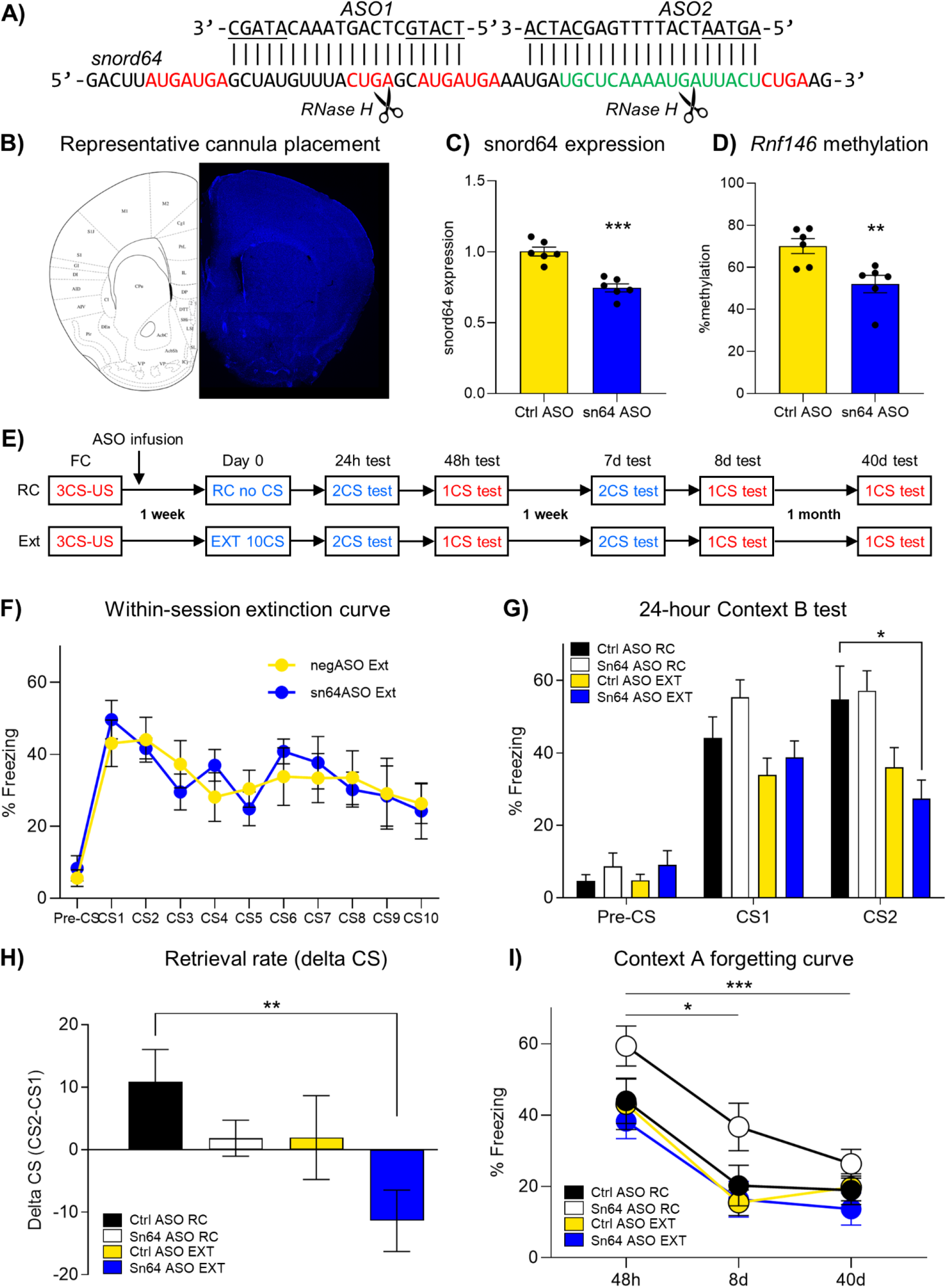
Knockdown of snord64 in the mouse prefrontal cortex stabilises fear memory and enhances fear memory retrieval. A) Two ASOs were designed against snord64 using a 5-10-5 gapmer design, with a total phosphorothioate backbone and edge nucleotides (underlined) modified with 2’-methoxyethyl.RNase H is expected to cut near the indicated locations within snord64. Red: C, C’, D’ and D motifs; green: *Rnf146* targeting element. B) Representative image of cannula track showing location of ASO infusion into the infralimbic PFC. C) 24 hours after injection of a cocktail of ASO1 and ASO2 into the mouse prefrontal cortex, expression of snord64 was reduced (Student’s t-test, n=6, p=0.0001.) D) In the same tissue samples, methylation of *Rnf146* was reduced (Student’s t-test, n=6, p=0.0079.) E) Timeline of procedures. FC = fear conditioning, RC = retention control, Ext = fear extinction. Red text indicates that a behavioural trial was performed in Context A, the context of fear training; blue text indicates alternative Context B. F) Within-session extinction curve showing 10 CS presentations. No difference was observed between snord64 knockdown and control mice (two-way ANOVA, n=9-10, no significant effect of group p=0.9089.) G) 24 hours after Ext or RC training, a 2-CS recall test in Context B showed a significant reduction in freezing during the second CS presentation for snord64 knockdown but not control mice (one-way ANOVA F(3,32)=4.9, p=0.0061, Dunnett’s post-hoc test ctrlASO-RC vs sn64ASO-Ext, p=0.0160.) H) Delta CS scores (CS2-CS1) show a significant difference in the rate of retrieval of fear extinction memory during the 24h test in Context B (one-way ANOVA, n=9-10, significant difference of means p=0.0263, Sidak’s multiple comparisons test p=0.0082.) I) Forgetting curve drawn from recall tests performed in Context A 48 hours, 8 days and 40 days after Ext/RC training shows a main effect of time with freezing scores showing a significant reduction from the highest levels overall at 48 hours to the lowest levels at 40 days (two-way repeated measures ANOVA, F(2,61)=68.33, p<0.0001) and a main effect of group with snord64 ASO treated RC mice showing the highest levels of freezing overall (F3,31=3.38, p=0.0306) suggesting attenuated forgetting in snord64 knockdown RC mice.

ASOs were infused bilaterally into the infralimbic region of the medial prefrontal cortex (ILPFC) through a surgically implanted guide cannula (Figure 6B). 24 hours after ASO infusion, a significant reduction in snord64 expression was observed in mice that had received the snord64-targeting ASOs (Figure 6C). In the same animals, a proportional reduction of *Rnf146* methylation was observed in the snord64 knockdown group, providing strong correlative evidence that snord64 is the methylation guide for *Rnf146* mRNA (Figure 6D).

To investigate the functional role of snord64 in fear-related memory, we first trained a cohort of ILPFC-cannulated mice to fear an auditory cue using a typical delay fear conditioning protocol. The mice were then infused with a cocktail of the two snord64-targeting ASOs or the negative control and, following a one-week delay, underwent weak extinction training consisting of 10 presentations of the CS, or a time-equivalent retention control. Following extinction training or non-reinforced exposure to a novel context, several tests of fear recall were performed to assess the influence of snord64 knockdown on memory (Figure 6E). No difference between groups was observed within-session for fear extinction (Figure 6F).

24 hours after extinction or retention control training, mice were tested with 2 presentations of the CS in Context B (the extinction context). There was no difference in the average freezing score between snord64 knockdown and control animals in either the retention control or extinction groups (Figure 6G). However, the response of extinction-trained snord64 knockdown mice to the two presentations of the cue differed from controls. Delta CS scores were calculated by subtracting the freezing score during CS1 from the freezing score during CS2, to quantify the difference in response to the first presentation of the CS relative to the second presentation within the test session (Figure 6H.) The average delta CS score for the mice which received the control ASO was positive, indicating increased freezing during the second presentation of the conditioned stimulus. For the snord64 knockdown mice, the delta CS scores were negative, showing that these animals froze more to the first than the second presentation of the cue. Based on these data, we conclude that snord64 knockdown increases the rate of memory retrieval which, in the context of fear extinction, may suggest an enhancement in processes related to memory updating.

We also observed a difference in the stability of the original fear memory over time in the animals that did not undergo fear extinction. During a recall test conducted in Context A (the context of fear conditioning) 48 hours after the extinction/retention control training day, retention control animals which received snord64-targeting ASOs showed an increase in freezing to the auditory cue compared to retention control animals which received negative control ASOs (Figure 6I). This effect persisted during a subsequent Context A test 8 days after weak extinction, but returned to equivalent baseline levels after 40 days. This finding indicates that a reduction in snord64 led to a strengthening of the original memory trace, resulting in attenuated forgetting over time, thereby suggesting that localized snord64 activity may constrain memory stability and prevent updating in the original training context.

## Discussion

Although 2’-O-methylation is increasingly recognised as an important internal modification of rRNA, relatively few publications have explored the possibility of snoRNA-guided mRNA methylation. Here we show that snoRNAs, typically assumed to have constitutive functions, also exhibit experience-dependent patterns of expression in the adult brain, and that the orphan snoRNA snord64 modulates memory processes and modifies the mRNA encoding a neuroprotective ubiquitin ligase, *Rnf146*.

We detected 84.7% of annotated mouse C/D box snoRNAs in the PFC. The library preparation method limited our ability to detect some unusual snoRNAs, such as those exceeding 110nt, as well as some RNAs with related functions such as ‘mini-snords’, C/D box scaRNAs or lncRNAs with snoRNA domains (Jorjani et al., 2016). The absence of some other snoRNAs cannot be explained by their size or structural features, and may represent selective expression. For instance, mice have four very similar copies of snord88, all of which target 28S:C3357; we observed only three in the PFC. In addition, a few snoRNAs with a unique target site were not detected. To determine whether these snoRNAs are truly absent from the brain, or were not detected by sequencing for structural or unknown reasons, the specific ribosomal positions they target could be assayed for the presence of 2’-O-methylation.

Another noteworthy finding is that 21.3% of C/D box snoRNAs expressed in the medial PFC are orphans, without complementarity to ribosomal RNA. Despite the lack of known targets for these RNAs, many are abundantly expressed and 31 were regulated dynamically in response to fear extinction training. Previous studies have reported that several orphan C/D box snoRNAs have brain-specific patterns of expression (Cavaillé et al., 2000; Runte et al., 2001; Rogelj, 2006). Most of these snoRNAs are associated with two imprinted genomic regions which give rise to neurological disorders when their appropriate imprinting or function is disrupted. The snord113/114 complex snoRNAs are encoded within the DLK1-DIO3 locus, dysregulated expression of which is associated with Temple and Kagami-Ogata syndromes (van der Werf et al., 2016). Snord64, snord115, snord116, and several other C/D box snoRNAs occur within the critical region for Prader-Willi syndrome, a disorder characterized by intellectual disability and neonatal growth deficiency followed by hyperphagia and obesity (Angulo et al., 2015). Prader-Willi syndrome is caused by loss of a paternally expressed imprinted domain on human chromosome 15, which includes multiple tandem repeats of both snord115 and snord116, both of which are implicated in the Prader-Willi syndrome phenotype (reviewed by Cavaillé, 2017). A previous study reported that snord115 is upregulated in the hippocampus in response to contextual fear conditioning (Rogelj et al., 2003). We did not observe significant differential expression of snord115 in the PFC in response to fear extinction learning; however, we did observe a significant upregulation of snord116. A weakness of the present study is that we did not separate reads mapping to distinct snord115 and snord116 genomic clusters; for this reason, we may have missed differential expression of only some clusters of these repetitive snoRNAs. Given the importance of snord116 in particular to the Prader-Willi syndrome phenotype (Duker et al., 2010; Bieth et al., 2015; Fontana et al., 2017), further investigation of the response of specific snord116 copies to learning experiences is warranted.

Snord115 represents the first example of a snoRNA with a mRNA target, having been shown to interact with the mRNA encoding the serotonin receptor *Htr2c* (Cavaillé et al., 2000; Vitali et al., 2005). This interaction alters both splicing and adenosine-to-inosine editing of this mRNA, and through both effects, snord115 promotes production of the fully functional 5HT2C protein as opposed to less functional isoforms, contributing to appetite dysregulation in mice, although the magnitude and importance of this effect is controversial (Vitali et al., 2005; Kishore and Stamm, 2006; Glatt-Deeley et al., 2010; Kishore et al., 2010; Raabe et al., 2019; Hebras et al., 2020). Importantly, while one study showed that snord115 methylates an artificial target RNA (Vitali et al., 2005), no publications claim to have detected 2’-O-methylation of endogenous *Htr2c*. In our study, we demonstrate that endogenous *Htr2c* within the mouse PFC is methylated, and the high stoichiometry observed for this mark suggests that its function may be constitutive rather than regulatory. In addition to its relevance to Prader-Willi syndrome, this finding validates *Htr2c* as a positive control for future studies of snoRNA-guided 2’-O-methylation of mRNA in the brain.

To date, two publications have reported methylation of messenger RNA by C/D box snoRNAs. Elliott *et al* revealed that snord32a and snord51 target a coding sequence of peroxidasin (*Pxdn*) mRNA, which is 2’-O-methylated (Elliott et al., 2019). They found that the methylation level decreases when the snoRNAs are knocked down, resulting in reduced *Pxdn* mRNA levels but increased PXDN protein. They also reported direct impairment of translation by the methylation of the *Pxdn* coding sequence. Recently, van Ingen *et al* found that the orphan snoRNA snord113-6 is required for fibroblast cell survival and methylates multiple mRNAs within the integrin pathway; inhibition of this snoRNA reduced mRNA methylation and increased degradation of the targeted mRNAs, with mixed effects on subsequent protein expression (van Ingen et al., 2022). Furthermore, several studies have reported widespread 2’-O-methylation of eukaryotic mRNAs (Dai et al., 2017; Hsu et al., 2019), although the origin and function of these methylation sites is largely unknown.

Together, these findings suggest a regulatory mechanism by which 2’-O-methylation increases the abundance and stability of mRNA, but may impair its translation, likely depending on the position of the modification within the mRNA. This is highly plausible based on the known ability of 2’-O-methylation to block degradation of RNA by nucleases (Lapham et al., 1997; Yu et al., 1997) and enhance the stability of many RNA structural states (Prusiner et al., 1974; Kawai et al., 1991; Abou Assi et al., 2020), as well as to directly inhibit decoding of mRNA by the ribosome (Choi et al., 2018; Elliott et al., 2019). The value of this mechanism in the context of local synaptic translation is obvious; enhancing RNA stability while reducing translation would provide a mechanism for maintaining regulated levels of local protein synthesis at synapses while slowing mRNA turnover to reduce the required investment in RNA trafficking. Although no proteins are currently known to remove 2’-O-methylation from RNA, we speculate that such proteins may exist, and that active demethylation of 2’-O-methylated mRNAs at synapses could permit surges of translation and degradation of previously dormant mRNAs in response to stimulus. Our observation of higher stoichiometry of methylation of *Rnf146* at the synapse is consistent with this hypothesis.

*Rnf146*, also known as dactylidin and Iduna, is an attractive candidate gene for memory-related processes for multiple reasons. RNF146 is an E3 ubiquitin ligase which specifically ubiquitinates poly-ADP-ribosylated (PARsylated) proteins, resulting in their degradation (Zhang et al., 2011; Shen et al., 2018). One protein that is ubiquitinated by RNF146 is PTEN (Li et al., 2015), which is a negative regulator of memory and a reduction in PTEN activity has previously been shown to promote the formation of fear extinction memory (Murphy et al., 2017; Song et al., 2018). Other RNF146 targets include AXIN1 and AXIN2, which act as negative regulators of Wnt signaling; by targeting them for removal, RNF146 is an activator of Wnt signaling in several contexts (Zhang et al., 2011; Shen et al., 2018). Wnt signaling has known roles in memory formation, especially in the hippocampus (reviewed by Oliva et al., 2013; Fortress and Frick, 2016) Neuroprotective functions of RNF146, also related to Wnt signaling, have been observed during glutamate excitotoxicity (Andrabi et al., 2011; Yang et al., 2017), stroke (Andrabi et al., 2011; Belayev et al., 2017) and oxidative stress (Belayev et al., 2017). *Rnf146* messenger RNA is predominantly localized at the synapse, as shown in this study (Figure 4F) and others (Glock et al., 2020). We propose that a reduction in *Rnf146* methylation by snord64 knockdown leads to enhanced local translation of RNF146 protein at the synapse, resulting in increased Wnt signalling and enhanced capacity for memory updating. The link between RNF146 and the memory enhancement observed in snord64 knockdown mice is speculative, and future studies will need to determine whether snord64 knockdown influences the translation of *Rnf146* or alters the abundance of RNF146 protein expression at the synapse.

Another important open question is the functional relevance of the presence of snoRNAs at synapses of the medial PFC. The increased abundance of both snord64 and methylated *Rnf146* at synapses following fear extinction learning implies that stimulus-responsive methylation of *Rnf146* may be occurring at synapses. However, the methyltransferase fibrillarin, which is required for snoRNA-guided mRNA methylation, is not known to occur outside the nucleolus, although our laboratory and others have observed the messenger RNA encoding this methyltransferase at synapses (Cajigas et al.; Glock et al., 2020). Given that most copies of snord64 are located in the nucleus, and that around 50% of copies of *Rnf146* from the non-synaptosomal fraction are methylated, we assume that some *Rnf146* methylation occurs in the nucleus or nucleolus. Selective transport of methylated *Rnf146* to synapses would explain the different stoichiometry of methylation observed between synaptic and non-synaptic compartments, but would not explain the presence or dynamic changes of snord64 at synapses. For this reason, we propose that some copies of *Rnf146* are methylated by snord64 at synapses of the mouse prefrontal cortex, and that this process is upregulated by fear extinction learning. Nonetheless, the finding that multiple snoRNAs are present in the synaptic compartment of the medial PFC, and that at least four of them undergo dynamic localization shifts in response to fear extinction learning, suggests the existence of a new class of regulatory RNA that is present at the synapse.

In summary, we have expanded the understanding of dynamic, experience-dependent snoRNA expression in the adult brain, and identified a messenger RNA target, *Rnf146*, for the orphan snoRNA, snord64. The level of *Rnf146* methylation increases during fear extinction learning, thereby demonstrating that 2’-O-methylation can act as a dynamic epitranscriptomic mark on mammalian mRNA. Knockdown of snord64 proportionally reduces *Rnf146* methylation and enhances the stability of fear memory and efficiency of memory updating, revealing a novel epitranscriptomic mechanism affecting process associated with fear-related memory.

## Acknowledgements

The authors gratefully acknowledge grant support from the NIH (R01MH109588, to TWB and RCS), the NHMRC (GNT1181359, to TWB) and the Prader-Willi Research Foundation of Australia (1424-PWRFA, to LJL). LJL and SUM were supported by postgraduate scholarships from the Westpac Scholars Trust. LJL is supported by a postdoctoral fellowship from The University of Queensland. We thank Ms Rowan Tweedale for helpful editing of the manuscript.

